# Recommendations For a Standardized Approach to Histopathologic Evaluation of Synovial Membrane in Murine Models of Experimental Osteoarthritis

**DOI:** 10.1101/2023.10.14.562259

**Authors:** Alia M Obeidat, Sung Yeon Kim, Kevin G Burt, Baofeng Hu, Jun Li, Shingo Ishihara, Rui Xiao, Rachel E Miller, Christopher Little, Anne-Marie Malfait, Carla R Scanzello

## Abstract

**Background:** Synovial pathology has been linked to osteoarthritis (OA) pain in patients. Microscopic grading systems for synovial changes in human OA have been described, but a standardized approach for murine models of OA is needed. We sought to develop a reproducible approach and set of minimum recommendations for synovial histopathology in mouse models of OA.

**Methods:** Coronal and sagittal sections from male mouse knee joints subjected to destabilization of medial meniscus (DMM) or partial meniscectomy (PMX) were collected as part of other studies. Stains included Hematoxylin and Eosin (H&E), Toluidine Blue (T-Blue) and Safranin O/Fast Green (Saf-O). Four blinded readers graded pathological features (hyperplasia, cellularity, and fibrosis) at specific anatomic locations in the medial and lateral compartments. Inter-reader reliability of each feature was determined.

**Results:** There was acceptable to very good agreement between raters. After DMM, increased hyperplasia and cellularity and a trend towards increased fibrosis were observed 6 weeks after DMM in the medial locations, and persisted up to 16 weeks. In the PMX model, cellularity and hyperplasia were evident in both medial and lateral compartments while fibrotic changes were largely seen on the medial side. Synovial changes were consistent from section to section in the mid-joint area mice. H&E, T-blue, and Saf-O stains resulted in comparable reliability.

**Conclusions:** To allow for a standard evaluation that can be implemented and compared across labs and studies, we recommend using 3 readers to evaluate a minimum set of 3 pathological features at standardized anatomic areas. Pre-defining areas to be scored, and reliability for each pathologic feature should be considered.

## INTRODUCTION

Osteoarthritis (OA) is a debilitating joint malady characterized by structural joint pathology including degeneration of articular cartilage and remodeling of subchondral bone. All joint tissues can be affected by this disease and increasingly attention has turned to the role of the synovial membrane, as synovial inflammation is recognized to be an active component of OA pathogenesis. Clinically, synovitis detected by modern imaging techniques has been associated with more severe symptoms and more rapid progression of disease in patients. Furthermore, synovial changes detected early in disease may predict incident disease, as they are observed in some patients before signs of OA cartilage degeneration are detectable (1–3). At the tissue level, the histological pattern of synovial pathology in OA patients is complex and variable, owing to the highly dynamic processes involved in the diseased joint environment. To understand this complexity better, many investigators now routinely evaluate synovial histopathology in preclinical studies using established murine models of OA (4–7).

Several semi-quantitative histopathology grading systems have been utilized by different investigative groups to assess the severity and variability of the mouse synovial response **(Suppl. Table 1)**. These range from simple systems evaluating a single pathologic feature of the synovium (*i.e.*, lining hyperplasia, (25)) or evaluating multiple features on a single scale (8), to more complex grading systems evaluating multiple pathologic features (9,10). Other investigators have adapted human synovial pathology grading schemes (11,12) for use in mouse models (13–15). These approaches have been. informative, and each has its merits, but a standardized approach has yet to be adopted. In addition, specific limitations of published systems remain to be addressed. First, the approach of extrapolating grading systems developed for evaluating synovium from patients with arthritis (11,12) to assessment of synovitis in OA rodent models (7,16) can be problematic. Not only are there differences in histopathology between humans and other animals (17–19), but these systems have often been developed to distinguish OA from more highly inflammatory forms of arthritis (*e.g*., rheumatoid arthritis, RA) and may not be sensitive enough to capture the range and variability of more subtle pathologic changes observed in many OA mouse models. Second, synovitis can be more focal and patchy in OA than in RA, and there is considerable anatomic variability in the normal appearance of the synovium as well as the location of synovitis in the knee joint (4,19). Yet, there is currently no standardized approach to defining anatomic regions for scoring synovitis to capture this variability in rodent models. Third, many approaches compute a summation of multiple cellular and tissue parameters (*i.e.*, hyperplasia, fibrosis, inflammation) to arrive at one composite score, an approach likely adapted from scoring methods to assess histopathologic cartilage damage in OA models. However, the synovial response in OA does not clearly progress in distinct phases representing progressive tissue damage, as cartilage degeneration does (20–22). Moreover, this approach fails to provide distinctions between pathologies that might result from different cellular or molecular mechanisms. For example, two synovial specimens may have the same “summed” synovitis score despite having markedly different degrees of hyperplasia, fibrosis, or infiltration of leukocytes. The loss of granularity through reliance only on summed scores to define “synovitis” may impede understanding of the synovial response in experimental OA. Finally, it remains unclear whether different commonly used histochemical stains perform better in accentuating certain hallmarks of synovitis (*e.g.*, inflammation, fibrosis). All of these concerns indicate that standardization of grading protocols should be considered.

Many approaches to targeting “synovitis” have been tried in preclinical OA models, with varying success and an appreciation that some aspects of the synovial response may be protective (23,21,24). Advances in this field will depend upon a better understanding of spatiotemporal variations in synovial responses, and how they relate to synovial function in various models as OA develops and progresses. To this end, reproducible histologic assessments of synovial pathology in murine models of OA that are easily implemented and address some of the limitations of current methods are a critical starting point for reliable comparison of results among different research groups. Accordingly, the goal of the current study was to synthesize the best practices and features of existing systems into a standardized approach and set of recommendations for routine histological evaluation of synovial changes in murine models of experimental OA.

## MATERIALS AND METHODS

### Knee joint sections

Coronal and sagittal sections from murine knee joints collected as part of other studies (19,25) were included in this comparative analysis. Total of 109 male mice were used in the study. All animal procedures had been approved by the Institutional Animal Care and Use Committee at either Rush University Medical Center (Chicago, IL), the University of Pennsylvania and the Corp. Michael J. Crescenz VA Medical Center (Philadelphia, PA), or Kolling Institute (Sydney, Australia). All sections included were from male C57BL/6 mice wild-type (WT) mice that had been housed with food and water *ad libitum* and kept on 12-hour light cycles. Sections from two surgical models of OA were evaluated: the destabilization of the medical meniscus (DMM) and the partial meniscectomy (PMX) models. **Table 1** summarizes the groups included in this analysis.

**Table 1:**
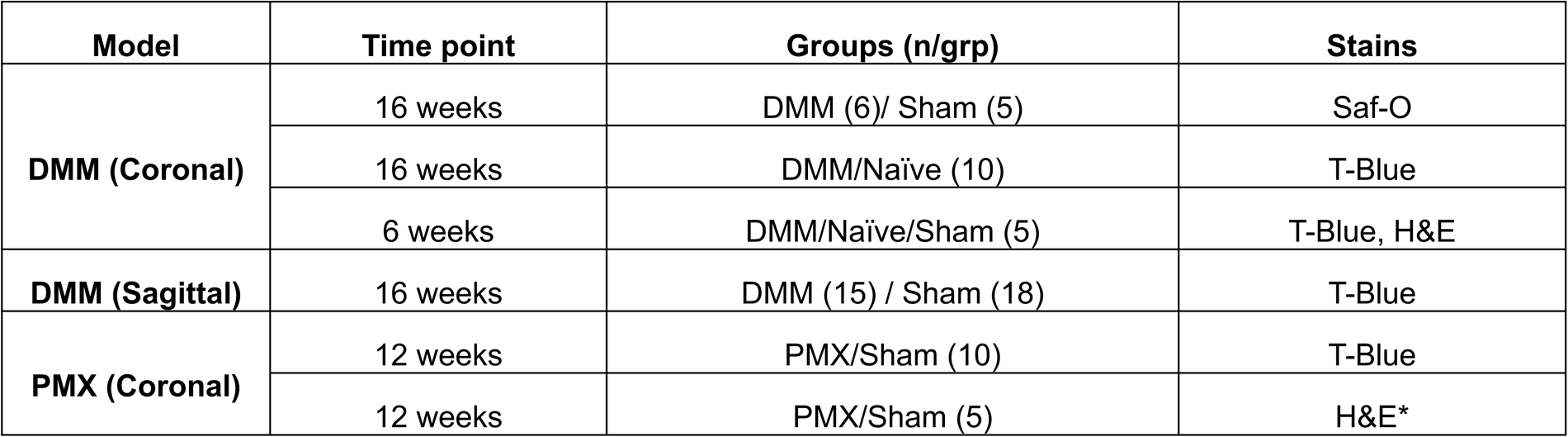
Summary of OA models, time points, sample size and stains included in the analysis. *4 sections per mouse evaluated in this group. All evaluations in other groups were done on a single midpoint section.

### OA model induction

#### Destabilization of the medial meniscus (DMM)

DMM (n=36) or sham (n=28) surgery was performed in the right knee of 10 to 12-week old male mice as described (4). Age matched naïve group (n=15) was added to the study. Briefly, the joint capsule was opened and the anterior medial meniscotibial ligament (MMTL) was cut. The knee was flushed with saline, and the incision closed. Sham surgery was identical to DMM surgery, except that the ligament was left intact. Mice were sacrificed at 6- or 16-weeks *post-*DMM or sham surgery.

#### Partial meniscectomy (PMX)

PMX (n=15) or sham (n=15) surgery was performed in the right knee of 10-week old male C57BL/6 mice, as previously described (5). All PMX surgeries were performed by the same surgeon. Briefly, mice were anesthetized by inhalation of isoflurane, and a medial parapatellar arthrotomy was performed. The MMTL was transected to release the anterior horn of the medial meniscus, and approximately 1/3-1/2 of the anterior portion of medial meniscus was cut. Mice were sacrificed 12 weeks after PMX or sham surgery.

#### Histology

Right knees had been previously collected, formalin fixed, and paraffin embedded. Knees were sectioned at 5 µm thickness, and mid-joint coronal or sagittal sections were stained using Toluidine Blue (T-Blue), Hematoxylin and Eosin (H&E), or Safranin-O/Fast Green (Saf-O) using routine methods. We evaluated the mid-joint area because cartilage pathology is most pronounced and consistent in this location in these models (33), and the synovial “gutters” (areas where the synovium reflects off the bone and onto the joint capsule; **Fig. 1A**) are easily identified in this area.

**Figure 1:**
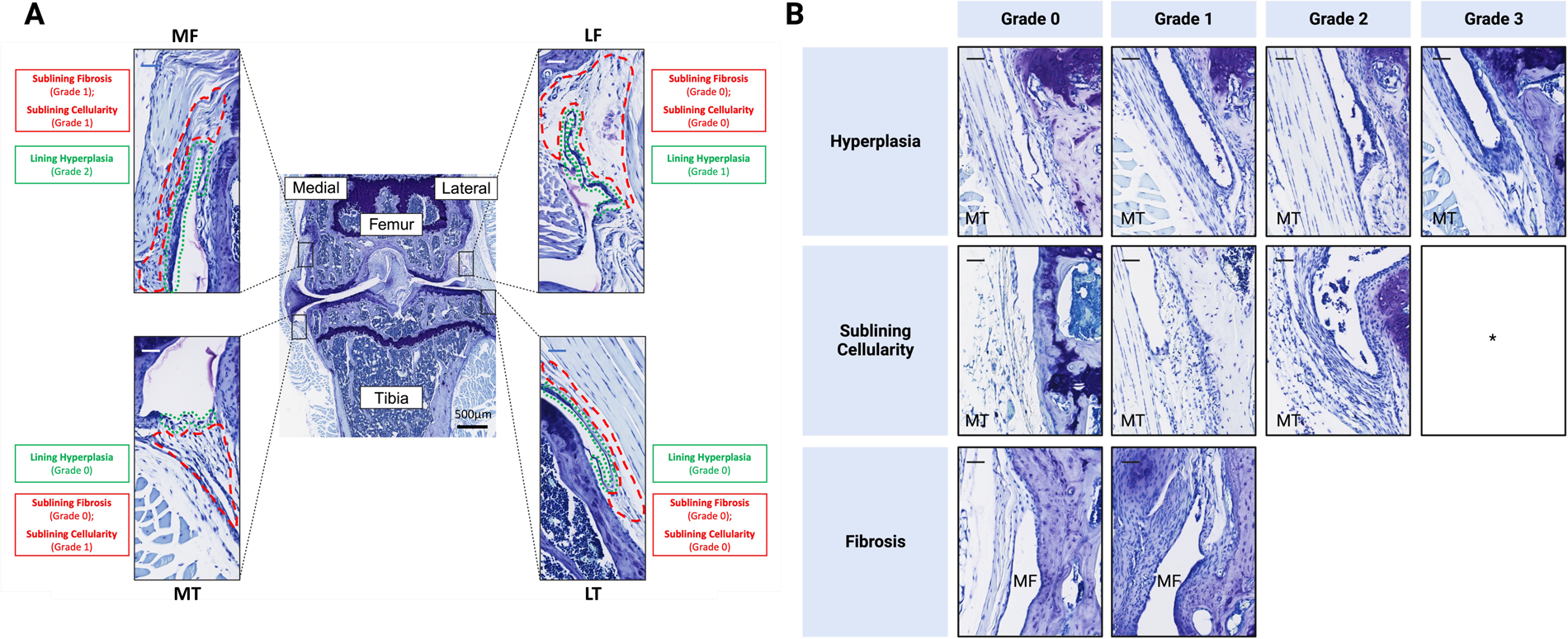
A) Four different anatomic locations of the synovium were assessed in coronal sections of murine knee joints: medial and lateral femur (MF and LF, respectively) and medial and lateral tibia (MT and LT, respectively). B) Representative histological images of each of the grades for three pathological features (hyperplasia, cellularity, and fibrosis). Scale bar= 50 µm. *Grade 3 sublining cellularity was not observed at the time points and experimental murine models of OA evaluated in this study. In some midpoint sections at MF gutter, no synovium could be identified where the capsule attached directly to the femur, even in uninjured/normal joints. In these cases, no score was recorded for this location.

#### Histopathologic Grading

Synovial pathology was assessed by 3-4 independent observers (AMO, SYK, BH, CRS, KGB). On coronal sections, we followed the approach reported by Lewis *et al* (30) and evaluated synovial changes at four regions: medial tibial (MT), medial femoral (MF), lateral tibial (LT) and lateral femoral (LF) gutters. (**Fig. 1A**). Synovium directly adjacent to the meniscus (pre-meniscal tissues or peri-meniscal plica) or directly attached to the bone was avoided, as described by Jackson *et al* (9). Four pathological features commonly included in published scoring methods **(Suppl. Table 1)** and typically found in OA models were assessed: lining hyperplasia, subintimal cellularity, sub-synovial inflammation, and subintimal fibrosis, according to (**Table 2** and **Fig. 1B**). Synovial hyperplasia refers to the thickness of the lining layer, cellularity is defined as cellular density of the synovial subintimal layer, and synovial fibrosis refers to the density of extracellular matrix staining in the synovial subintimal layer. As routine histologic stains used in this study do not specifically stain fibrotic changes to the matrix in sufficient detail, fibrosis was simply scored as absent (0) or present (1). Many published grading schemes include a category for “inflammation”, which is defined as subsynovial infiltration of mononuclear cells either in a perivascular or diffuse pattern (9,26). In our experience, routine histologic stains are insufficient to specifically identify whether murine subsynovial cellular infiltrates seen in OA models truly represent inflammatory cell types, and the definition of inflammation used here overlaps the description of cellularity. Thus, a separate inflammation score was also assessed to compare to cellularity scores. We tested multiple mid-joint section 12 weeks after PMX or sham surgery to determine whether multiple sections are needed to capture the synovial reaction versus using a single section. On sagittal sections, the same features were evaluated along the same scale, but the standard anatomic areas evaluated were the posterior femoral and tibial synovium (PFS and PTS, respectively) and the anterior femoral and tibial synovium (AFS and ATS, respectively) as shown in **(Suppl. Fig. 1).**

**Table 2:**
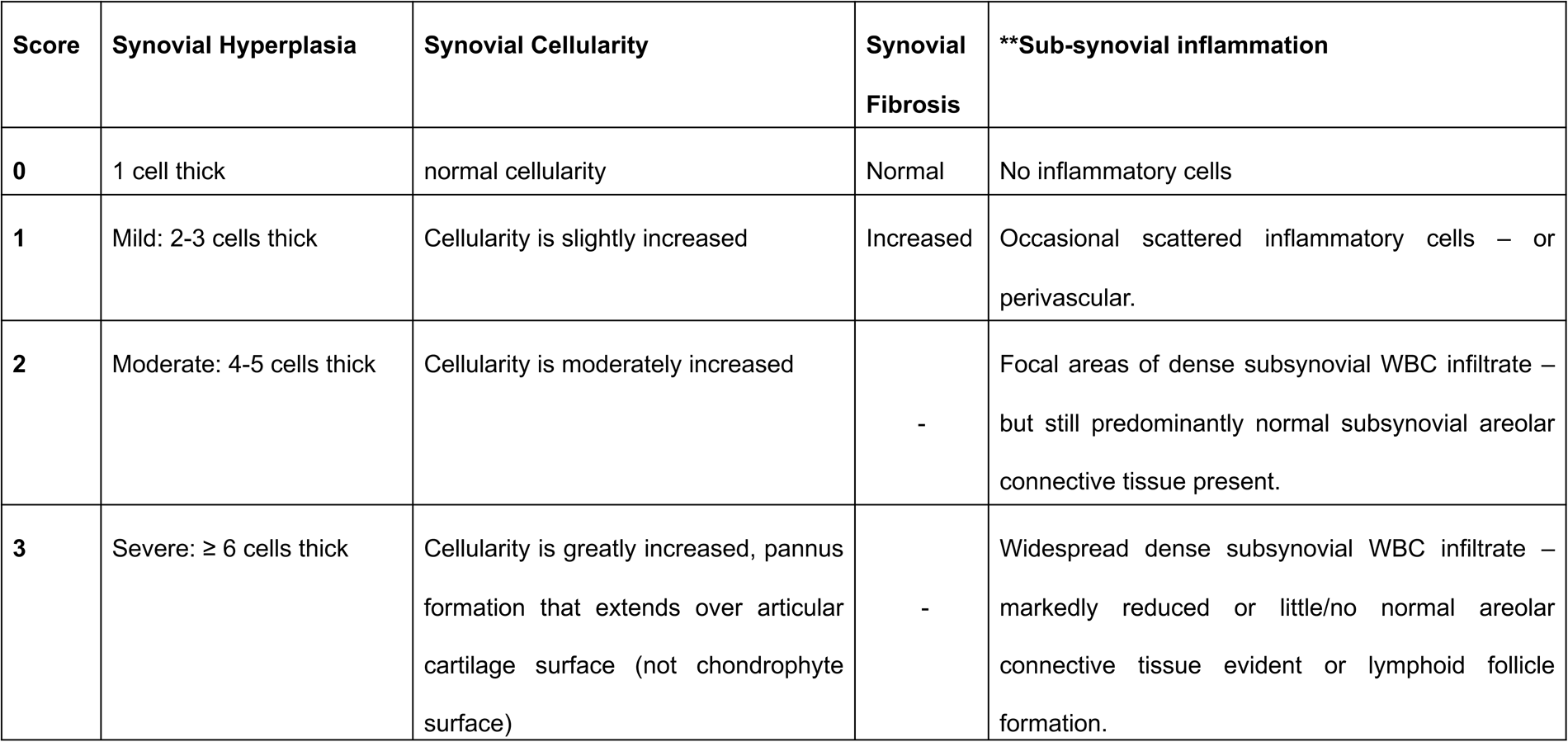
Synovial scoring description. ** Included for comparison to cellularity score.

During scoring sessions, one reader outlined the area to be evaluated at 20X magnification on digital images, so that all readers were assessing the same anatomic region (illustrated in **Fig. 1**). In order to determine a score, we stipulated that the degree of change needed to be present in 50% or more of the area being evaluated (*i.e.*, grade 2 hyperplasia needed to be present in at least half of the lining evaluated in a particular location). After evaluating a set of training slides together, all four scorers assessed the same images at the same time, were blinded to treatment groups, and graded independently and without discussion. Representative histological images of hyperplasia, cellularity, and fibrosis are shown in (**Fig. 1B**).

### Statistical Analysis

To assess the inter-rater reliability (IRR), we used Kendall’s coefficient of concordance (W) with adjustment for ties for the ordinal data (the variables graded from 0-3) and Fleiss’ Kappa tests (κ) for fibrosis which had a binary rating (0 or 1). The values for these reliability indices range between 0 and 1, with higher values for better agreement between readers. Though interpretation of reliability scores may vary from group to group, we classified scores from 0.41 to 0.60 as a moderate agreement, and 0.61 to 1.0 as a substantial agreement. Anything below 0.41 was determined to be a poor and unacceptable agreement. Once reliability was determined, the mean scores of all 4 readers were reported. Normality of data was assessed using the D’Agostino & Pearson test. Non-parametric Kruskal-Wallis was used to compare multiple groups and joint compartments followed by Dunn’s post-hoc. Non-parametric Mann-Whitney test was employed for two-group comparison, using the Holm-Šidák method to adjust for multiple comparison. Statistical analyses were performed using RStudio and GraphPad Prism 9.4.1. All data are presented as mean±95%CI.

## RESULTS

### Inter-rater reliability of each variable and the effect of stain on grading

Four blinded observers evaluated the synovial changes in two mouse models of OA. In order to measure the degree of agreement between the different readers, we assessed the inter-reader reliability (IRR) using Kendall’s coefficient of concordance (W) for hyperplasia and cellularity, and Fleiss Kappa (κ) for fibrosis. Using knee sections from mice 12 weeks after PMX or sham surgery stained with T-blue or H&E, we found acceptable moderate to substantial IRR using either stain to detect lining hyperplasia, sub-lining cellularity, and fibrosis (**Table 3**). In general, inflammation scores had slightly lower, but still acceptable, agreement between different readers across T-blue and H&E stains.

**Table 3:**
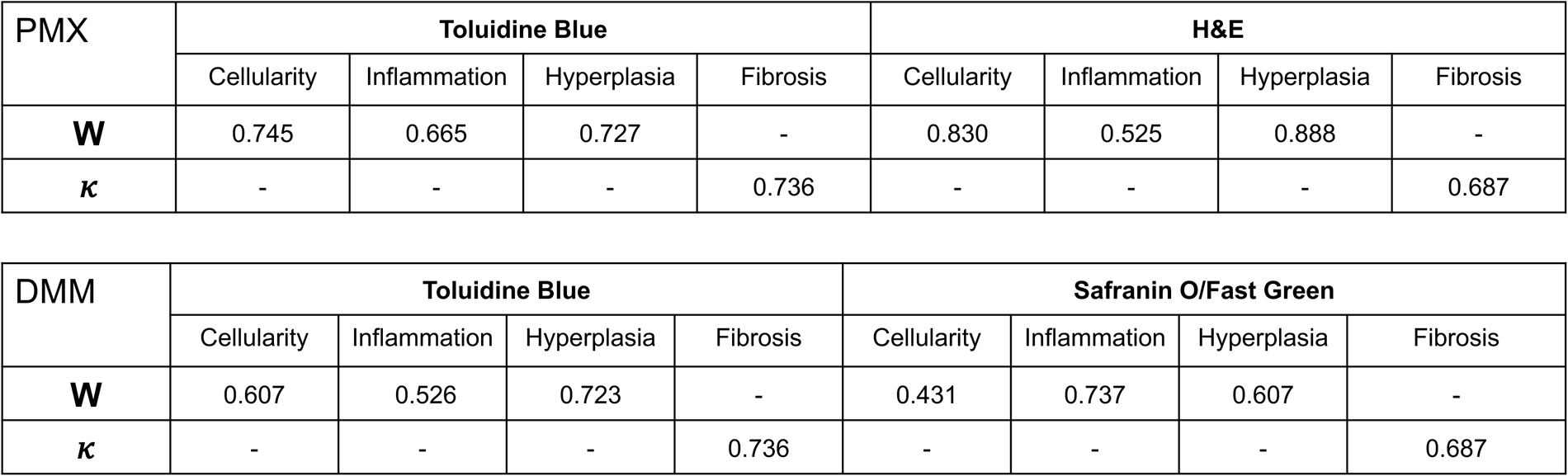
Inter-reader reliability scores of murine knee joints 12 weeks after PMX or sham surgery and 16 weeks after DMM or sham surgery. Kendall’s coefficient of concordance (W), for the variables graded from 0-3, and Fleiss Kappa tests (κ) for fibrosis which was graded 0 or 1.

To further explore the effect of other commonly used stains on synovial scoring, we repeated this reliability analysis using T-blue and Saf-O stained knee sections from mice 16 weeks after DMM surgery and age-matched naïve mice. Here again, both stains showed good reliability among readers in all 3 parameters (**Table 3**). Cellularity assessed on Saf-O stained sections showed the lowest reliability in our hands but still indicated moderate agreement (W=0.43). Given the previously mentioned overlap between “cellularity” and “inflammation”, and the inability of routine histologic stains to definitively identify inflammatory cell types, the more straightforward cellularity category was considered sufficient on its own and separate inflammation scores were not included for all subsequent analyses.

### Anatomic variation across four anatomic areas

Due to the known anatomic variability of synovial architecture, we compared synovial changes at the 4 predefined anatomical areas on coronal sections (medial tibial, medial femoral, lateral tibial and lateral femoral gutters) (**Fig. 1A**). Knee sections from mice 6 and 16 weeks after DMM or sham (n=5/group, n=10/group respectively) and 12 weeks after PMX or sham surgery (n=5/group) were compared.

#### DMM model

6 weeks *post*-DMM, we observed increased lining hyperplasia and sub-lining cellularity in the medial tibial and medial femoral gutters compared to age matched sham and unoperated controls (**Fig. 2A,B**). Changes were very mild (mean±SD, medial tibial and medial femoral hyperplasia naïve = 0.15±0.22 and 0, sham = 0.22±0.27 and 0.25±0.25, DMM = 0.75±0.3 and 0.75 ±0.5 respectively, cellularity naïve = 0.2±0.11 and 0, sham = 0.4±.0.28 and 0.35±.0.13, DMM = 0.75±0.3 and 1.05±0.75 respectively). We also detected trends toward increased fibrosis in the medial tibial and femoral gutters (**Fig. 2C**) at this time point, albeit statistically not significant. In this model, no significant synovial pathology compared to controls was observed on the lateral side (**Fig. 2A-C**), but of note there was slightly greater hyperplasia and cellularity even in controls on the lateral side compared with the medial locations, potentially indicating some natural anatomic variation in the thickness and cellularity of the synovium even in naïve joints. At 16 weeks *post*-DMM, increases in hyperplasia compared to age matched sham mice were no longer observed at the medial locations, but increased cellularity and fibrosis persisted (**Fig. 2D-F**). Synovial cellularity and fibrosis were not significantly different between the 6- and 16-week time points *post*-DMM (cellularity for MT DMM 6 weeks *vs.* 16 weeks p=0.18, fibrosis p=0.88).

**Figure 2:**
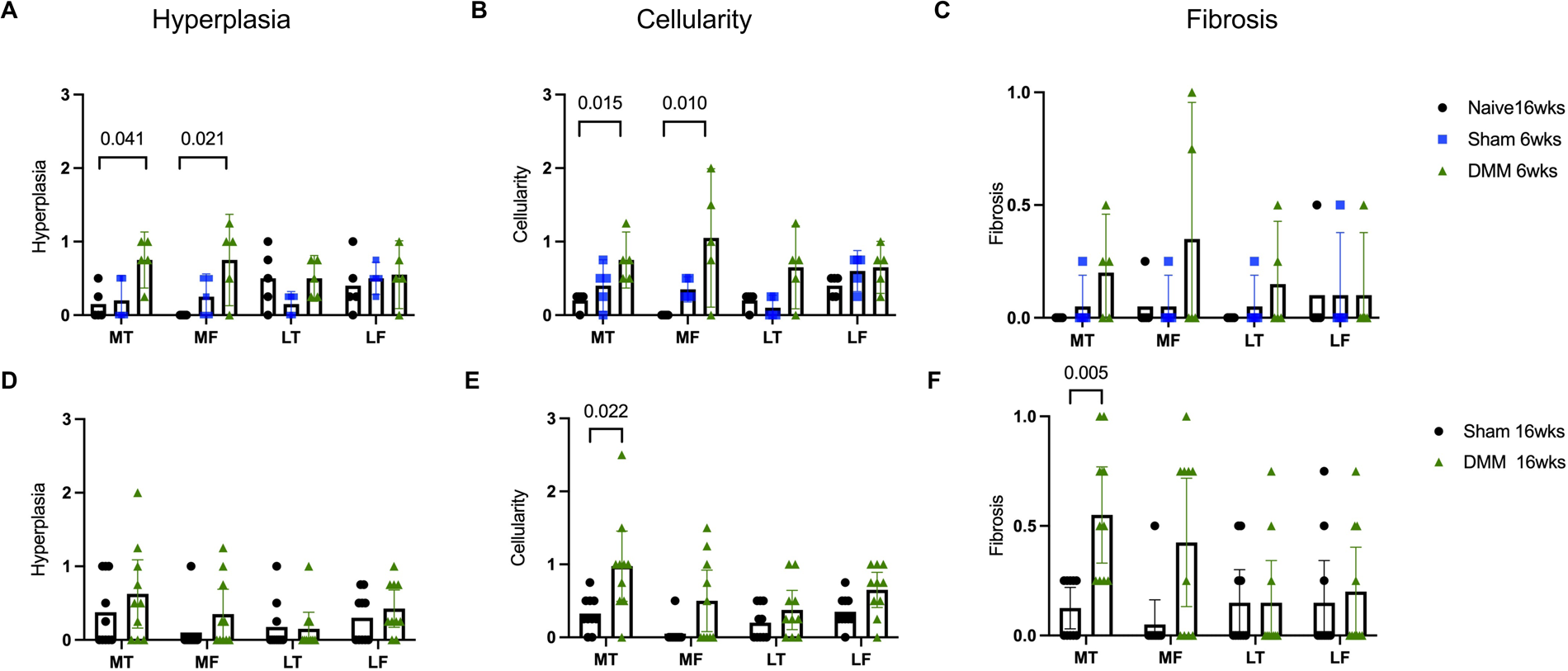
(A-F) Histopathologic scoring of synovial hyperplasia, cellularity and fibrosis for medial tibial (MT), medial femoral (MF), lateral tibial (LT) and lateral femoral (LF) areas in 16-week old naïve mice and mice 6 weeks after sham or DMM surgery (n=5/group) in (A-C); and mice 16-week after sham or DMM surgery (n=10/group) in (D-F). Each score represents the average score of 4 blinded scorers. (A-C) Nonparametric Kruskal-Wallis with Dunn’s posthoc. (D-F) Multiple unpaired Mann-Whitney tests corrected for with Holm- Šidák method. Mean ± 95% CI.

#### PMX model

We next evaluated synovial histopathology using this method in knee sections from mice taken 12 weeks after PMX surgery. As shown in (**Fig. 3A**), we observed a significant increase in hyperplasia in the medial compartment of PMX operated mice compared to sham mice. Increases in subsynovial cellularity were observed in all four anatomic locations in this model, although with more pronounced changes on the medial side of the knee (**Fig. 3B**). Synovial fibrosis, like hyperplasia, was significantly elevated primarily on the medial side in PMX operated *vs*. sham-operated joints. In general, synovial pathologic features were slightly higher in this model compared to the DMM model at both timepoints, although differences were not statistically significant.

**Figure 3:**
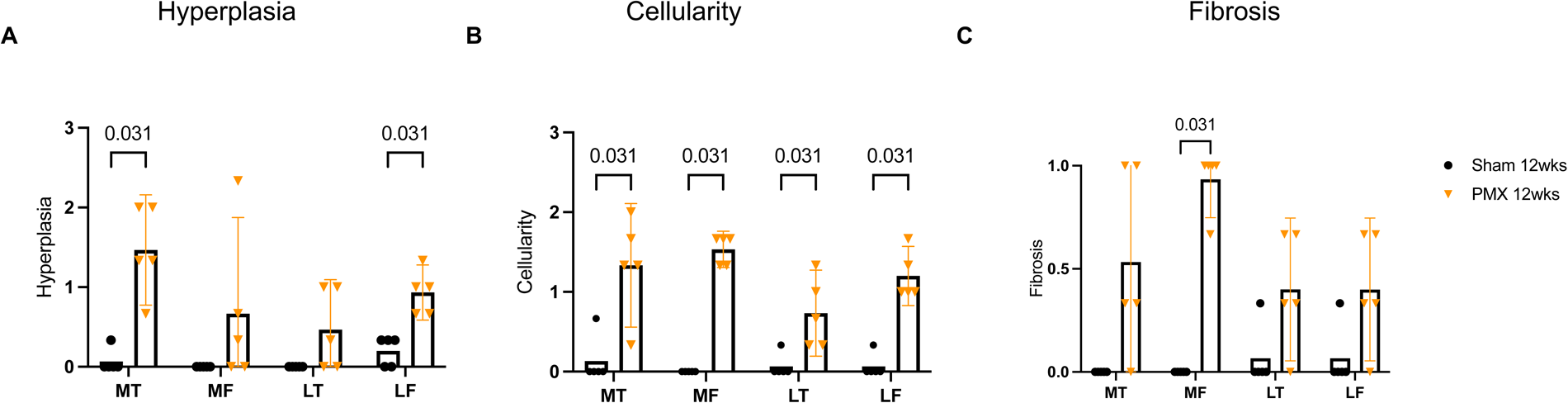
(A-C) Histopathologic scoring of synovial hyperplasia, cellularity and fibrosis respectively for medial tibial (MT), medial femoral (MF), lateral tibial (LT) and lateral femoral (LF) areas in mice 12 weeks after PMX or sham surgery (n=5/group). Each score represents the average score of 4 blinded scorers. Multiple unpaired Mann-Whitney tests corrected with Holm-Šidák test. Mean ± 95% CI.

### Using single versus multiple sections

In the routine evaluation of cartilage histopathologic damage, many scoring systems suggest the use of multiple sections throughout the joint (8), although this practice may not always be necessary and a single mid-point section may be sufficient (27). To decide whether multiple sections are needed to capture the synovial reaction in these models, we compared scoring using multiple coronal sections versus a single mid-joint section. For this analysis, we used 4 H&E-stained sections per mouse 20μm apart spanning the mid-joint area (mid-joint area as described in (28)). For each of the 3 parameters (synovial hyperplasia, cellularity, and fibrosis), we first summed individual readers’ scores from all four joint areas for each knee section, to get a single score for the whole joint for each reader. Then, the summed score for each feature was averaged for the 4 readers. The average scores for each of the 4 sections per mouse knee were then compared within the same treatment group in (**Fig. 4**). Whole joint scores for synovial hyperplasia, cellularity, and fibrosis were very consistent and did not vary widely from section to section (**Fig. 4A-C**).

**Figure 4:**
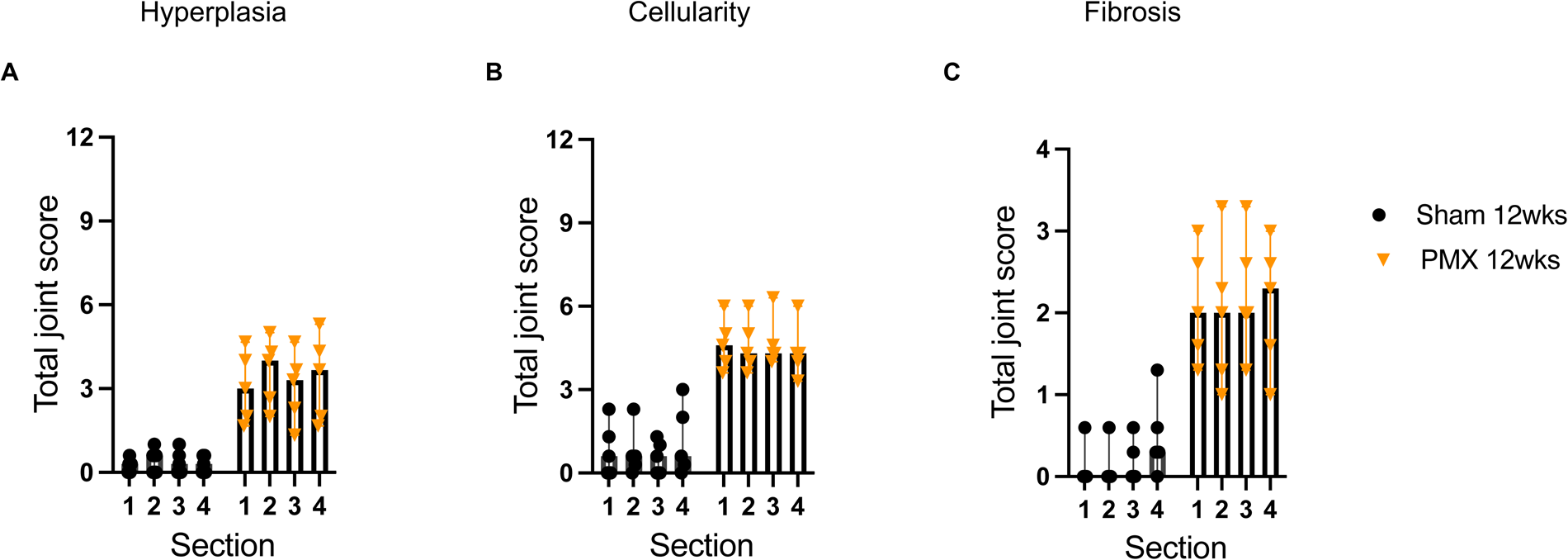
(A-C) Histopathologic scores of synovial hyperplasia, cellularity and fibrosis respectively summed for 4 joint areas in mice 12-week after sham and PMX surgery (n=5/group). 4 mid-joint sections per mouse were used to evaluate synovial changes, sections were H&E stained. 4 sets of data are shown in x-axis, each set represents one section per mouse for n=5 mice/group. Kruskal-Wallis test with Dunn’s post-hoc. Mean ± 95% CI.

### Use of sagittal sections

Across studies, the choice of section orientation varies by research objective. Although coronal sections are used most commonly, sagittal sections may be preferred or necessary for certain studies. To demonstrate application of this scoring method into the sagittal plane, three independent graders scored mid-sagittal sections from DMM (n=15) and Sham (n=18) mice 16 weeks *post*-surgery, grading four standard regions as depicted in **(Suppl. Fig. 1)**. If the research objective requires evaluation of the synovial lining of the infrapatellar fat pad (IFP), we further suggest evaluation of this area on sections from the intercondylar notch where the IFP is most clearly visualized (in addition to the other four locations). Scoring categories remain the same (synovial hyperplasia, cellularity, fibrosis), and representative examples are shown in **(Suppl. Fig. 1)**. No significant differences between sham and DMM groups (16 weeks post-surgery) was observed in synovial histopathology within the sagittal section PFS region **(Suppl.** **Fig.1B**). We assessed IRR using previously mentioned methods **(Suppl.** **Fig.1C**).

## DISCUSSION

Synovial pathology in OA has attracted much attention since reports that linked synovial inflammation and effusions to joint pain (29–31). Several scoring systems have been applied to the evaluation of synovial changes in human OA (11,12,32,33), and several approaches to assess histopathology in mouse models have been reported (summarized in **(Suppl. Table 1)**. There is, however, no consensus on a standardized approach that can be easily implemented. In addition, many systems combine several features of synovial pathology into a single summed “synovitis” score. Although this “summing” method is useful for quantifying progressive cartilage histopathologic damage of OA, the pathological changes in the synovium do not necessarily reflect progressive degenerative changes. Instead, synovial changes appear to be more dynamic and reactive to changes in the inflammatory, degenerative, and traumatic joint environment (reviewed in (34)). Rather than quantifying progressive tissue damage, the goal of a synovial histopathology assessment should be to capture the severity, fluctuation, and anatomic variation of different pathological features observed in commonly used murine models. These issues highlight the need for a standardized approach to describe synovial pathology in murine models of OA. In addition, we sought a method that would allow enough granularity to distinguish different processes affecting the lining layer and the sublining layer separately, but still be easily implemented by most research groups to allow standardization even when a Veterinary Pathologist is not available. Our recommendations, which were informed by currently published methods, are summarized in (**Table 4**). Of note, we have utilized this system in a previous studies (35,36) with reproducible results to gain insight into aging-related changes to the synovium in mice.

**Table 4:**
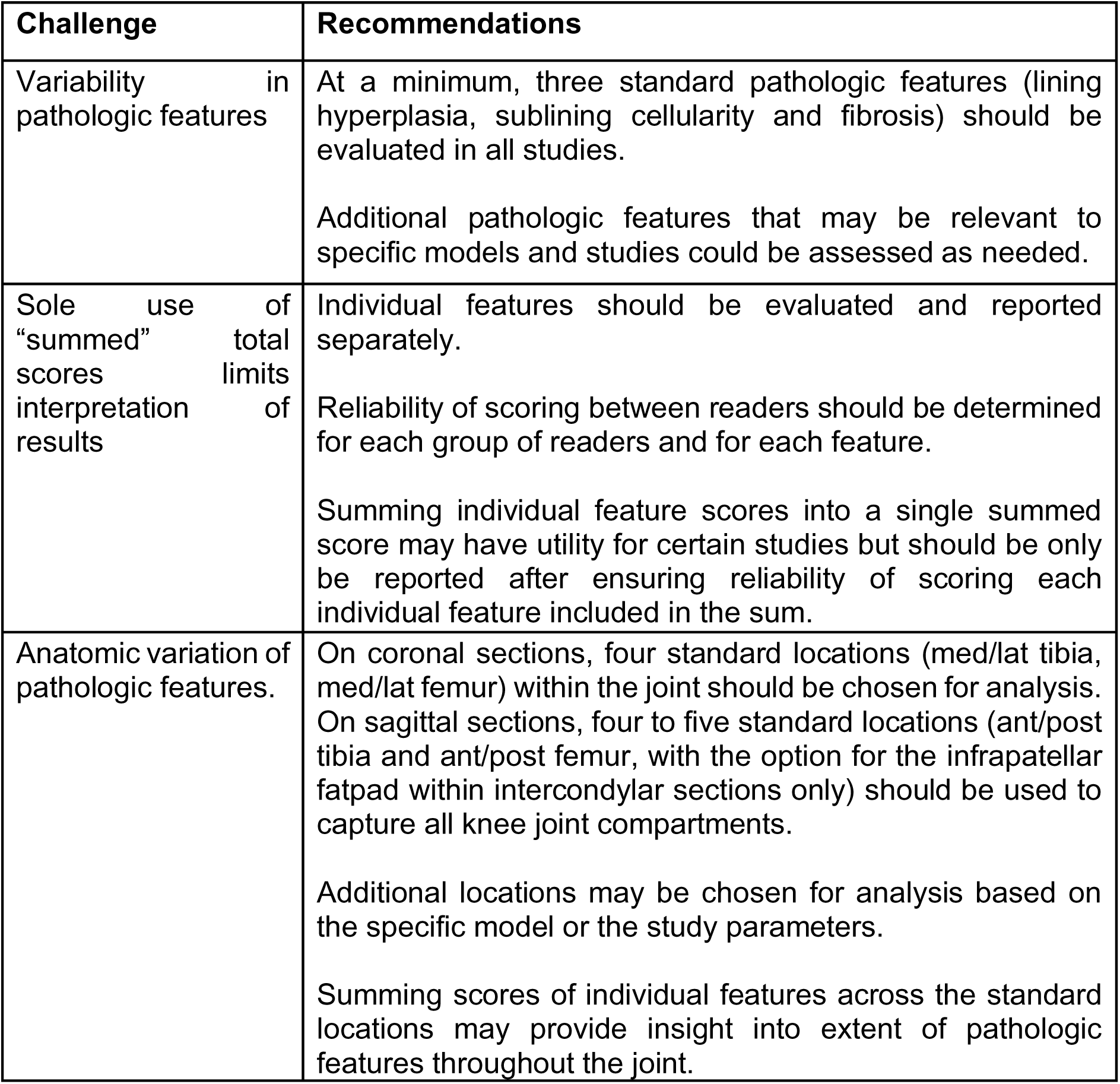
Recommendations for synovial histopathologic evaluation.

First, to capture the different patterns of pathologic change that affect the lining and sublining regions, we suggest that a minimum of three individual features be assessed separately. These include lining hyperplasia, sublining cellularity and sublining fibrosis. Although many scores include a separate category of “inflammation”, we suggest that the level of detail provided by common histopathologic stains is likely insufficient to definitively determine whether the increased cellularity in many murine models represents leukocytic inflammation or fibroblastic expansion, unless evaluated by a trained pathologist. Thus, for this approach the simpler category of “cellularity” should capture any inflammatory cell infiltration, and is highly reliable across graders. If increased subsynovial cellularity is observed, more specific techniques such as immunostaining and cytometry should be used to accurately distinguish inflammatory cell populations in the murine synovium.

Second, we recommend reporting individual features separately (**Table 2**), to allow for a more granular understanding of how specific features of the synovial reaction evolve with time in different models, and how these separate features impact joint health and disease. There may be utility in calculating a summed “synovial pathology” score for individual studies (14), but this should only be done after confirming and reporting the reliability of scoring the individual features that are included in the sum. Summed scores can be translationally useful and roughly comparable to human MRI grading of “synovial inflammation”, which captures global synovial abnormality. .

Third, to capture the anatomic variation in the synovial reaction we suggest assessing four standardized locations throughout the joint on coronal sections (4-5 on sagittal sections) to capture all synovial compartments of the knee joint. Locations suggested on coronal and sagittal sections are listed in (**Table 2** and depicted in **Fig. 1** **and Suppl. Fig. 1)**. In individual studies, it could be useful to calculate a “total joint” sum for each feature separately (hyperplasia, cellularity or fibrosis) to determine the extent of changes (how widespread features are throughout the joint). Finally, to address subjectivity of scoring we recommend a minimum of 3 readers for which a measure of agreement (IRR) should be performed. If there is high reliability between readers, an average score can be reported. Reliability within groups of scorers can be improved by doing “training” or practice sessions prior to blinded scoring.

We applied this method of evaluation of synovial change in two surgical mouse models of PTOA, the DMM and the PMX model, using coronal sections. Assessing the four standardized anatomic areas we found that medial tibial and medial femoral areas showed more pronounced changes compared to the lateral side in these models, consistent with other studies and the distribution of cartilage damage in these models (10,35). The medial predominance of synovial pathology was most pronounced in the DMM model (**Fig. 2 and 3**). These 2 models are induced by either surgical cutting of the MMTL or both the MMTL and medial meniscus respectively, which likely explains the predominantly medial side changes. In comparison, mild age associated changes in the synovium were previously observed in both medial and lateral joint compartments (35,36). These standard locations should first be chosen to allow comparison with other studies. The choice of additional locations to evaluate depends on the study, the model used, and the plane of sections (coronal versus sagittal plane).

Our analysis of sagittal sections showed adequate reliability, but in contrast to the coronal analysis no significant differences were observed between DMM and sham at 16- week timepoint. This can be attributed to the overall very mild changes observed in this model, and in addition the synovial locations evaluated on sagittal sections differ and are anatomically further from the mid-section of the joint where most cartilage pathology is observed in this model. However, the current study was not designed to specifically address differences in results obtained between coronal and sagittal sections in this model, and experimental variability including the use of different surgeons may contribute. Moreover, we only scored sections from mid-joint region, and did not evaluate sections at anterior and posterior locations in coronal sections or axial and abaxial locations in sagittal sections.

As histologic stain and number of sections per mouse evaluated often varies in reported studies **(Suppl. Table 1)**, we looked at the effect of both these variables. Knee sections using three of the most conventionally used stains for OA joint histopathological assessment (H&E, Saf-O and T-blue stains) were compared. H&E and T-blue stains showed slightly higher agreement between our readers compared to Saf-O stain, but all stains showed adequate reliability. Although the effect of stain was not very large, reliability should be optimized for a chosen stain before reporting results. To determine if the synovial reaction varied greatly from section-to-section, we evaluated four coronal sections per mouse from the PMX model. Hyperplasia, cellularity, or fibrosis exhibited minimal variability from section-to-section. Therefore, a single mid-joint section in this model was enough to reflect the severity of the synovial pathology, consistent with another recent study that showed that a single mid-sagittal section was sufficient to evaluate synovial pathology in common knee OA models (37).

The histopathologic assessment described in this study is considered an initial evaluation to guide subsequent analyses of the synovial reaction in models of OA. Recommendations are for a standardized, minimum data set for reporting histopathologic changes that is reliable and can be easily implemented without requiring a trained veterinary pathologist. A minimum set of anatomic locations and pathologic features (synovial hyperplasia, cellularity, and fibrosis) should be reported to allow for comparison across labs and models. Use of more detailed methods (9,37,38) and additional locations can be added to address individual study goals and model variability. Sections in the coronal plane offer the best summary of articular cartilage across the joint, capturing medial and lateral, tibial, and femoral surfaces, and in addition capture the medial and lateral synovial gutters. However, in studies evaluating changes to the synovial lining of the IFP, or in which the anterior and posterior compartments are of interest, the method can be adapted to sagittal sections to offer the most comprehensive view.

There are several limitations to the scoring system described, and histopathologic assessment in general, that deserve mention. In general, histopathologic grading is not very sensitive to subtle changes, particularly when evaluating the low-grade changes observed in many OA models, therefore further analyses using more sensitive methods should be considered. Specifically, detecting inflammatory cells and fibrosis in mouse models of OA is challenging using conventional stains, therefore additional specific cellular or matrix stains are recommended o fully evaluate these changes as needed. Additionally, it is recommended to have a minimum of 3 blinded readers to address subjectivity, and training done to optimize reliability, which requires time and labor. However, as a starting point a standardized system should be included whenever characterizing new murine models or testing therapeutics, as the synovial reaction is relevant to OA symptoms and overall health of the joint and the pathologic features that are most sensitive to change could provide insights into mechanisms of action.

## Conclusions

The current study suggests recommendations for a reproducible and easily implemented method to evaluate synovial pathology. The proposed method captures the main features of OA pathology in the lining and sublining regions and is applicable to common murine models of OA. This will allow for comparison of results across different research groups, therapeutic interventions, and animal models of OA. Evaluation of these three features will provide insight into whether changes are happening to the cellular (hyperplasia and celluarity) and/or the matrix (fibrosis) component of the synovium. Choosing standard anatomic locations as indicated will provide insight into where changes are localized in specific models. Application of this method should provide a reproducible data set and starting point for more detailed studies as needed to address individual study objectives.

## Contributions

Conception and design: AMO, CRS, REM, CBL, AMM.

Analysis and interpretation of the data: AMO, SYK, KGB, BH, RX, CRS

Drafting of the article: AMO, SYK, CRS

Critical revision of the article for important intellectual content: All authors

Final approval of the article: All authors

Statistical expertise: RX

Obtaining of funding: CRS, AMM, CBL, REM, AMO.

Collection and assembly of data: AMO, SYK, KGB, BH, SI, JL, CRS

## Suppl. Fig. legends

**Suppl. Table 1:**
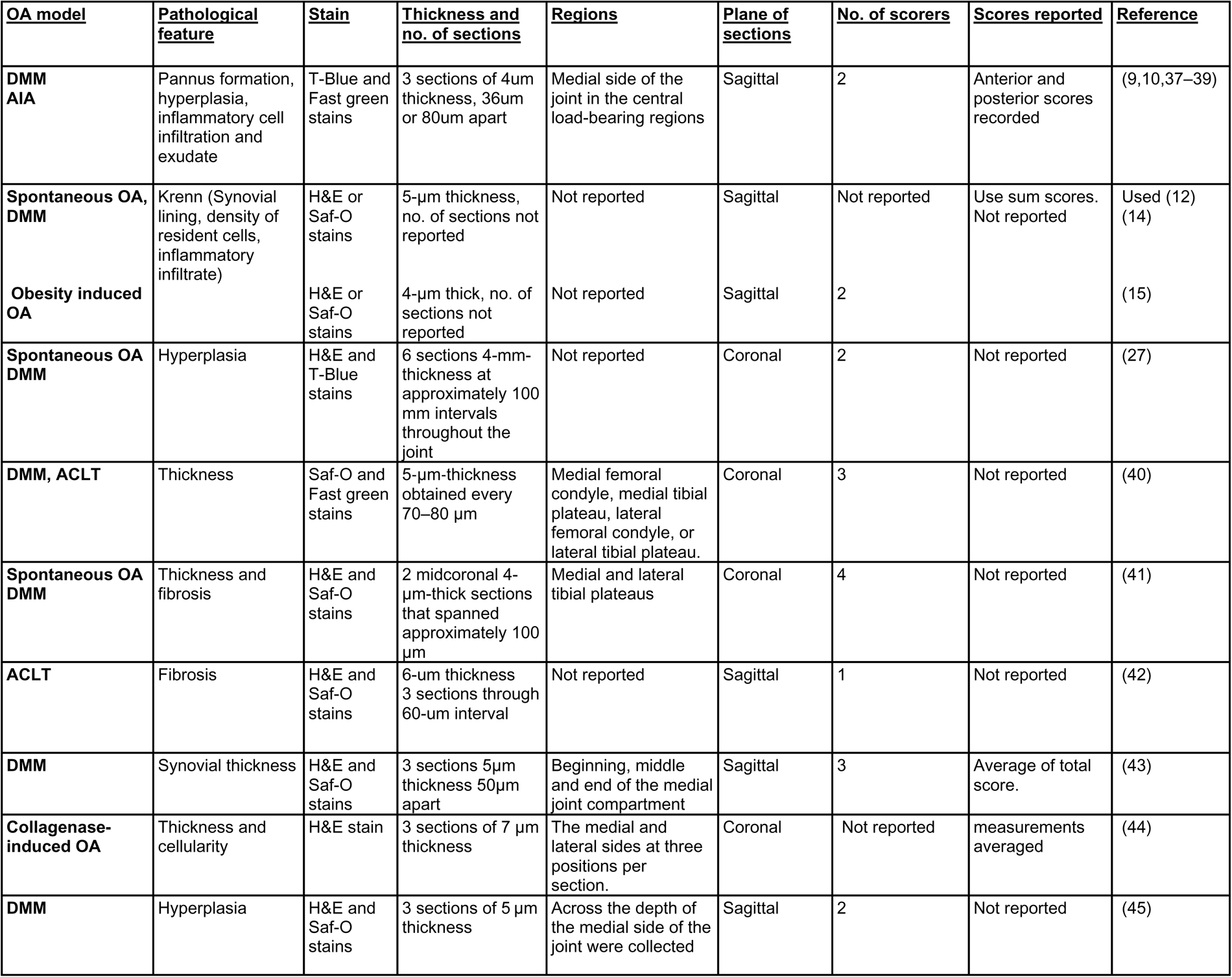
Summary of grading systems used for evaluation of synovial pathology in mouse models of OA.

**Suppl. Fig. 1:**
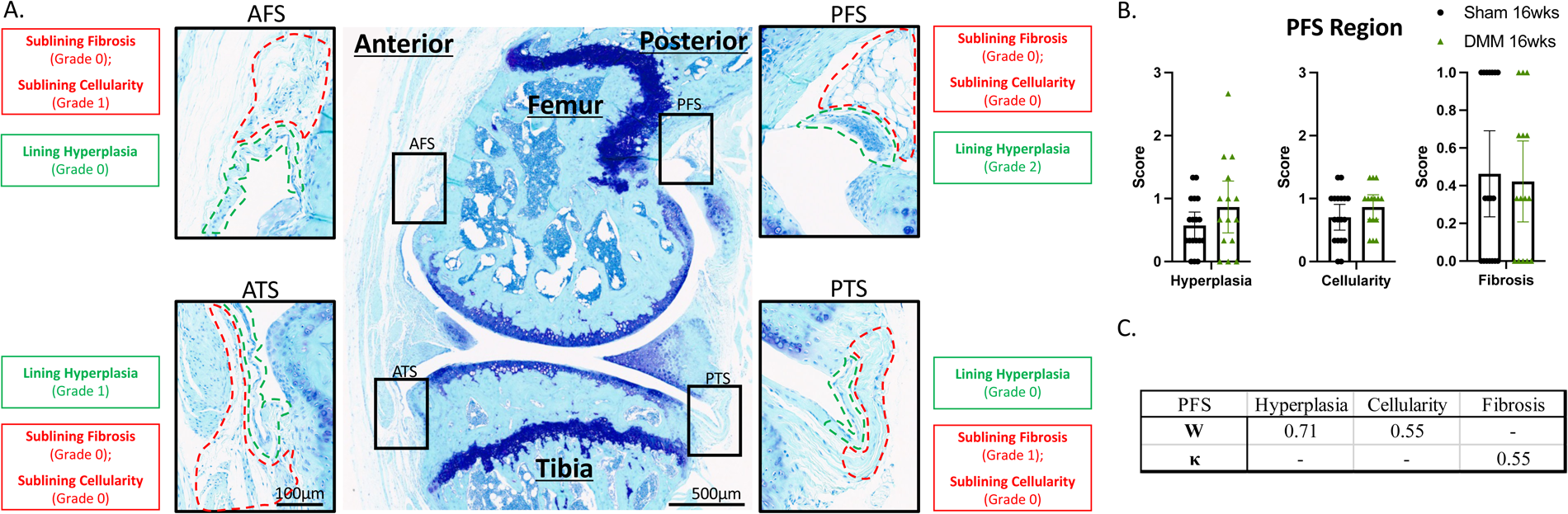
A) Four different anatomic locations of the synovium was assessed in T- Blue stained sagittal sections of murine knee joints: posterior femoral and tibial synovium (PFS and PTS, respectively), anterior femoral and tibial synovium (AFS and ATS, respectively). Green dashed line: lining layer. Red dashed line: sublining layer. Scale bars as indicated. B) Histopathologic scores of synovial hyperplasia, cellularity and fibrosis respectively in mice 16-week after sham (n=18) and DMM (n=15). C) Inter-reader reliability scores using Kendall’s coefficient of concordance (W), for the variables graded from 0-3, Hyperplasia and Cellularity, and Fleiss Kappa tests (κ) for Fibrosis which was graded 0 or 1.

